# Cell- and state-specific plasticity of striatal glutamatergic synapses is critical to the expression of levodopa-induced dyskinesia

**DOI:** 10.1101/2024.06.14.599055

**Authors:** Weixing Shen, Shenyu Zhai, Veronica Francardo, Qiaoling Cui, Zhong Xie, Tatiana Tkatch, M. Angela Cenci, D. James Surmeier

## Abstract

Levodopa-induced dyskinesia (LID) is a debilitating complication of symptomatic therapy in Parkinson’s disease (PD). Although there is compelling evidence that striatal pathophysiology is a major driver of LID, the circuit-specific mechanisms contributing to dysfunction remain obscure. This lack of clarity is reflected in the limited options for diminishing established LID. To address this gap, molecular, cellular, and behavioral strategies were used to interrogate striatal indirect pathway spiny projection neurons (iSPNs) in a mouse model of LID. These studies revealed that LID induction led to an up-regulation of GluN2B-containing N-methyl-d-aspartate receptors (NMDARs) specifically at iSPN glutamatergic synapses. This up-regulation was correlated with increased numbers of ‘silent’ glutamatergic synapses in the hours after levodopa treatment. In this ‘off-state’, long-term potentiation (LTP) of iSPN glutamatergic synapses was readily induced and this induction was blocked by antagonists of adenosine type 2 receptors (A2aRs) or GluN2B-containing NMDARs. Systemic administration of the A2aR antagonist tozedenant at the beginning of the off-state significantly reduced the development of LID. More importantly, specifically knocking down the expression of *GRIN2B* mRNA in iSPNs dramatically attenuated both development and expression of LID, without compromising the beneficial effects of levodopa on movement. Taken together, these studies demonstrate that dyskinesiogenic doses of levodopa trigger cell-specific synaptic adaptations during the off-state that make an important contribution to the network pathophysiology underlying LID and suggest that targeting GluN2B-containing NMDARs in iSPNs could be therapeutically useful.

## Introduction

Parkinson’s disease (PD) is the second most common neurodegenerative disorder ^1, 2^. The cardinal motor symptoms of PD – bradykinesia and rigidity or tremor – are caused by loss of functional dopaminergic neurons in the substantia nigra pars compacta (SNc). In the healthy brain, the release of dopamine (DA) by SNc neurons modulates basal ganglia circuitry, promoting movement vigor and goal-directed actions ^3^. In the early stages of PD, when a substantial population of SNc dopaminergic neurons remain, boosting their release of DA by administration of its blood-brain barrier penetrant precursor (L-3,4-dihydroxyphenylalanine or levodopa) effectively alleviates motor symptoms. However, as the disease progresses, higher oral doses of levodopa are required to achieve symptomatic benefit and brain concentrations of DA become dysregulated – rising to abnormally high levels for hours and then falling back to very low levels as systemic levodopa wains. These fluctuations lead to uncontrolled, dyskinetic movements when DA levels are high (on-state) and to severely impaired movement when they fall (off-state) ^4, 5^.

The mechanisms underlying levodopa-induced dyskinesia (LID) are poorly understood and, as a consequence, therapeutic strategies for diminishing their severity are limited ^6^. Several lines of evidence point to the importance of aberrant striatal synaptic plasticity in the induction and expression of LID ^5, 7–9^. In the healthy brain, transient alterations in striatal DA signaling in response to action outcomes modulate the induction of corticostriatal synaptic plasticity, altering the probability that the action will be repeated in the future ^10–12^. In late-stage PD patients given high doses of levodopa, not only is the relationship between DA and action outcome severed, alterations in DA concentration are sustained for hours. Indeed, during the on-state, when striatal DA levels are high, aberrant enhancement of intrinsic excitability and long-term potentiation (LTP) of corticostriatal synapses on D1 DA receptor (D1R)-expressing, direct pathway spiny projection neurons (dSPNs) appears to play a key role in triggering dyskinetic movements ^9, 13–15^. Consistent with this view, chemogenetic or optogenetic stimulation of dSPNs mimics LID ^4, 14, 16^.

Although dSPNs undoubtedly make a significant contribution to the network pathophysiology underlying LID, there are compelling reasons to think that aberrant plasticity in D2 DA receptor (D2R)-expressing indirect pathway SPNs (iSPNs) also contributes ^9, 16^. Normally, iSPNs are thought to promote context appropriate action by suppressing inappropriate, competing actions ^17^. This learned activity is widely thought to depend upon DA-driven adjustments in the strength of iSPN corticostriatal glutamatergic synapses. Unlike the situation in dSPNs, DA signaling (associated with reward) triggers long-term depression (LTD) of active iSPN corticostriatal synapses, presumably lessening their ability to suppress appropriate actions. In contrast, dips in DA signaling associated with negative outcomes disinhibits iSPN adenosine 2a receptors (A2aRs), promoting LTP of active corticostriatal synapses, which presumably leads to a ‘veto’ of that action in the future ^11, 18, 19^. This precisely timed, action-outcome-linked sculpting of iSPN synapses is not restored by levodopa treatment in late-stage PD patients lacking SNc dopaminergic neurons. Rather, these mechanisms appear to be hijacked for hours following levodopa treatment, driving iSPN synaptic plasticity in one direction during the on-state (down) and then in the opposite direction (up) during the off-state – all independently of behavioral outcomes.

Precisely how these alternating periods of plasticity remodel iSPNs is unclear. Several reports have highlighted the potential importance of alterations in N-methyl-d-aspartate receptors (NMDARs) at striatal glutamatergic synapses in LID ^20–22^. As at other glutamatergic synapses, NMDARs are necessary for the induction of LTP at corticostriatal synapses ^11, 23,24^. However, it is uncertain whether any of these changes are happening specifically in iSPNs and, if so, whether they impact dyskinetic behavior. A clue that these synaptic processes could be important is that dyskinesiogenic doses of levodopa trigger the restoration of iSPN axospinous glutamatergic synapses, which are lost following DA depletion ^13, 15, 25^. Using a combination of electrophysiological, genetic, pharmacological, and behavioral approaches, the studies described here show that dyskinesiogenic doses of levodopa up-regulate GluN2B-containing NMDARs in iSPNs, but not in dSPNs. This up-regulation leads to a significant elevation in the abundance of ‘silent’ glutamatergic synapses in the off-state and the induction of A2aR-dependent LTP. Connecting these cellular changes to behavior, A2aR antagonism in the off-state reduced the induction of LID. In addition, selectively suppressing expression of the GluN2B subunit in iSPNs significantly diminished not only the induction of LID, but the expression of established LID. Taken together, these studies provide a fundamental insight into mechanisms underlying LID and point to novel therapeutic strategies to alleviate them.

## Results

### LID induction was correlated with an up-regulation of GluN2B-containing NMDARs in iSPNs

Although there is compelling evidence that alterations in striatal glutamatergic signaling contribute to LID, the molecular and cellular specificity of these changes are poorly defined ^4, 5^. To help fill this gap, electrophysiological and pharmacological methods were used to assess the relative contribution of α-amino-3-hydroxy-5-methyl-4-isoxazolepropionic acid receptors (AMPARs) and NMDARs to iSPN glutamatergic, excitatory postsynaptic currents (EPSCs) in *ex vivo* brain slices from naïve, 6-hydroxydopamine (6-OHDA) lesioned, and 6-OHDA-lesioned mice induced to express dyskinesia by repeated administration of levodopa. Briefly, *Drd2*-eGFP bacterial artificial chromosome (BAC) transgenic mice were subjected to unilateral medial forebrain bundle (MFB) injections of 6-OHDA (Fig. 1a). Three to four weeks later, the extent of the lesions was assessed using a forelimb-use asymmetry test (also called cylinder test) ^9, 26, 27^. Mice with a robust lesion [>85% striatal tyrosine hydroxylase (TH) loss] ^16, 28^ were given either a short (3–5 days) or long (12–14 day) regimen of levodopa at a dose inducing dyskinesia in all animals (3 mg/kg/day, i.p.) ^16^ (Fig. 1b). Mice were then sacrificed, striatal brain slices prepared and dorsolateral striatal (DLS) iSPNs patch clamped in the presence of the GABA_A_ receptor antagonist gabazine (10µM). The relative contribution of AMPARs and NMDARs to electrically evoked EPSCs was estimated by measuring current amplitudes at −70 mV and then +40 mV in the presence of an AMPAR antagonist NBQX (10 µM) (Fig. 1c, d). In iSPNs from untreated, 6-OHDA lesioned mice, the NMDAR/AMPAR ratio was elevated (Fig. 1c, d); this shift was attributable to a reduction in AMPAR abundance ^13, 15, 25^. The elevation in the NMDAR/AMPAR ratio was also evident in iSPNs from lesioned mice treated with dyskinesiogenic doses for either short (3-5 days) or long periods (12-14 days) (Fig. 1c, d). However, NMDAR currents in the levodopa-treated iSPNs decayed significantly slower than those in untreated iSPNs (Fig. 1c, e).

**Figure 1:**
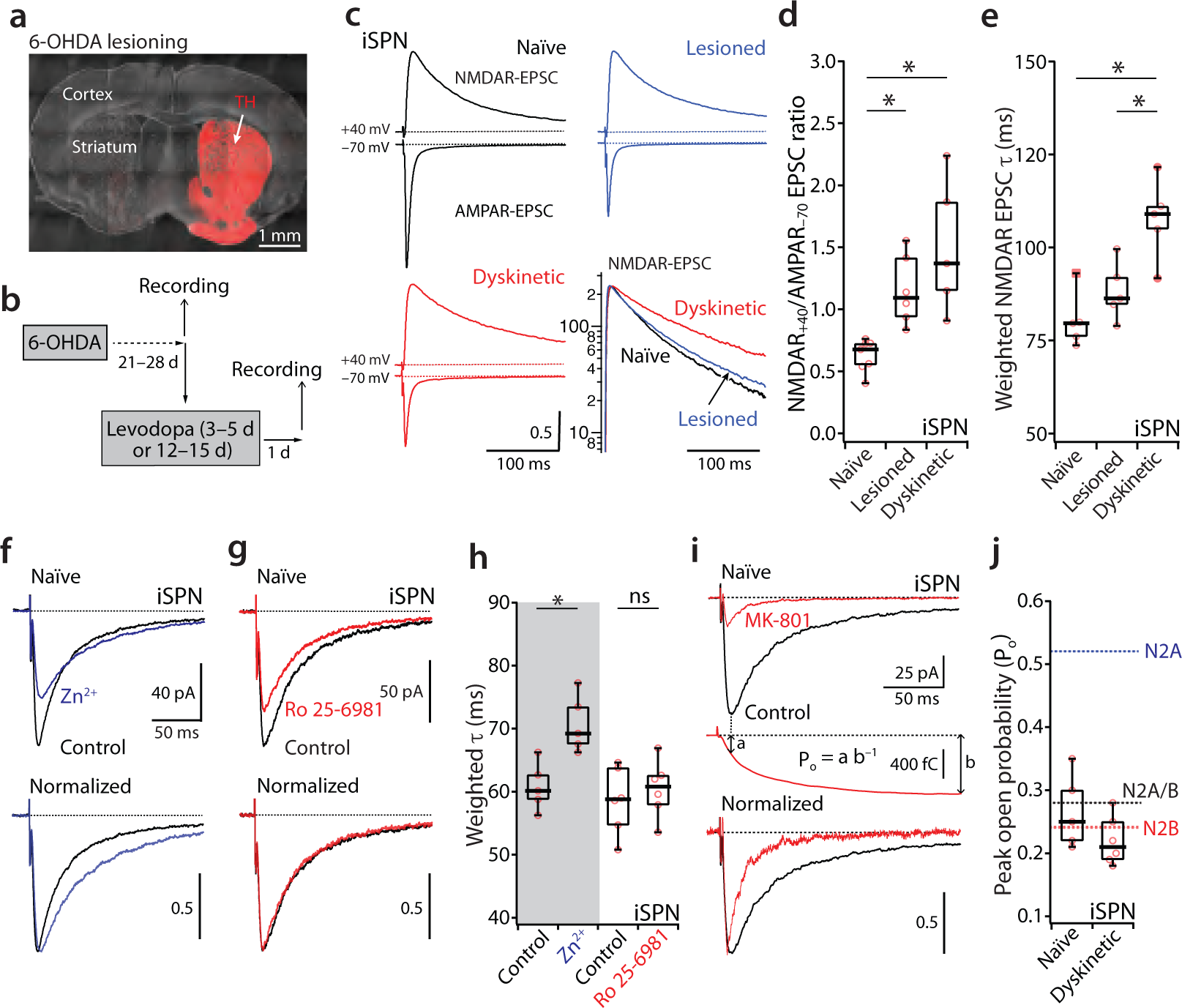
Up-regulation of iSPN GluN2B-containing NMDARs after LID induction. **(a)** Light microscopic image of a coronal section illustrating the loss of immunoreactivity for tyrosine hydroxylase (TH, red) after the unilateral MFB 6-OHDA lesioning. (**b**) Experimental timeline. (**c**) Averaged EPSC traces recorded at –70 mV (AMPAR-EPSC) and then +40 mV in the presence of the AMPAR antagonist NBQX (10 µM, NMDAR-EPSC) from a representative naïve, lesioned or dyskinetic iSPN. EPSC traces are normalized to the same height of NMDAR-EPSCs. The decay of these NMDAR currents was typically biexponential, as shown in the semi-logarithm plot at the lower right. The NMDAR current in the levodopa-treated iSPN decayed significantly slower than untreated iSPNs. (**d**) Box plot summary of the ratio of the NMDAR EPSC (measured at the holding potential of +40 mV in the presence of 10 µM NBQX) to the AMPAR EPSC (measured at –70 mV) in iSPNs from naïve (n = 7, from 4 mice), lesioned (n = 6, from 4 mice) and dyskinetic (n = 5, from 4 mice) states. Box plot boxes indicate upper and lower quartiles; whiskers specify upper and lower 90%. * p<0.05, Mann-Whitney rank sum test. (**e**) Box plot summary of the weighted iSPN NMDAR EPSC decay time constants from naïve, lesioned and dyskinetic states. * p < 0.05, Mann-Whitney test. (**f**) Bath application of GluN2A selective antagonist Zn^2+^ (100 µM with 10 mM tricine) decreased the size of the NMDAR currents and prolonged its de-activation. (**g**) Bath application of the GluN2B selective antagonist Ro 25-6981 (1 µM) also decreased the size of the NMDAR currents but did not change their de-activation kinetics. (**h**) Box plot sum of alterations of iSPN NMDAR current decay kinetics by the selective GluN2A antagonist (n = 5, from 3 mice, p < 0.05, Wilcoxon signed rank test) and the selective GluN2B antagonist (n = 6, from 4 mice, p > 0.05, Wilcoxon test). * p < 0.05; ns not significant. (**i**) NMDAR EPSCs from a naïve iSPN in control (black trace) and in 25 µM MK-801 (red trace) showing the reduction of the peak amplitude and the acceleration of the de-activation and the charge transfer (middle). In MK-801 the ratio of charge at the time of the EPSC peak (a) to the total charge (b) gives an estimate of the probability of channels having opened by the time of the peak. (**j**) In this plot, peak open probabilities (P_o_) of GluN2A-type NMDARs (0.52, N2A, blue dashed line), GluN2B-type NMDARs (0.24, N2B, red dashed line) and GluN2A/B-type NMDARs (0.28, N2A/B, black dashed line) are from the study of hippocampal synapses by Tovar et al ^36^, in which the subunit-specific P_o_s were obtained by using GluN2A or 2B knockout mice. The peak open probabilities of naïve (n = 5, from 3 mice) and dyskinetic (n = 6, from 3 mice) iSPNs fell near the value GluN1/N2A/B or GluN1/N2B receptors.

NMDARs are tetramers, consisting of two GluN1 subunits and two GluN2 subunits ^29–31^. In adult forebrain neurons, the majority of NMDARs are di- and tri-heteromers of GluN2A and/or 2B subunits ^29, 32–39^. NMDARs with GluN2B subunits deactivate more slowly than those with only GluN2A subunits ^36, 40–42^, suggesting that the slowing of NMDAR currents following levodopa treatment could be a manifestation of an increasing contribution of GluN2B-containing NMDARs to synaptic currents.

To test this hypothesis, pharmacological tools were used to assess the relative contribution of GluN2A- and GluN2B-containing NMDARs to evoked currents in iSPNs. In *ex vivo* brain slices from naïve mice, bath application of the GluN2A-prefering inhibitor Zn^2+^ (100 µM with 10 mM tricine) reduced the amplitude of NMDAR-mediated currents and slowed their de-activation ^36, 43^ (Fig. 1f, h). Bath application of the GluN2B inhibitor Ro 25-6981 (1 µM) also reduced the amplitude of the NMDAR currents but did not alter their de-activation kinetics (Fig. 1g, h). This observation is consistent with work in hippocampal neurons showing that the GluN2A subunit determines NMDAR channel de-activation kinetics ^33, 35–37^.

To gain more insight into the contribution of GluN2A and GluN2B subunits to synaptic NMDARs, the peak open probability (P_o_) was estimated using the high-affinity NMDAR open channel blocker MK-801 (25 µM) ^36, 44^. To estimate P_o_, the ratio of charge transfer at the peak of the EPSC (a) to the total charge transfer (b) was computed in the presence of MK-801 (P_o_=a/b) (Fig. 1i) ^36^. Because GluN2A-containing NMDARs activate more rapidly than GluN2B-containing ones, the P_o_ measured at the peak of the EPSC is correlated with subunit composition, being near 0.5 for di-heteromeric GluN1/GluN2A channels and roughly half that for di-heteromeric GluN1/GluN2B channels (Fig. 1j) ^36^. Furthermore, the P_o_s of tri-heteromeric GluN1/GluN2A/2B channels is similar to that of di-heteromeric GluN1/GluN2B receptors, indicating that GluN2B subunit determines NMDAR channel opening kinetics ^36^. The P_o_s of iSPN NMDARs from naïve and dyskinetic animals fell near the value of di-heteromeric GluN1/GluN2B or tri-heteromeric GluN1/GluN2A/B receptors (Fig. 1i, j), suggesting that synaptic NMDARs in iSPNs were largely GluN2B-containing tri- or di-heteromeric receptors.

To provide another assessment of the contribution of GluN2B-containing channels to synaptic NMDARs in different states, the ability of the GluN2B-selective antagonist Ro 25-6981 (1 µM) to reduce currents was examined in *ex vivo* brain slices from naïve, 6-OHDA-lesioned and dyskinetic mice. Both *Drd1*-tdTomato and *Drd2*-eGFP mice were used in these experiments to allow the cellular specificity of effects to be determined. In dSPNs, the contribution of GluN2B-containing NMDARs to the synaptic currents was similar in each of the three conditions (Fig. 2a, b, c). This was not the case for iSPN, however. In iSPNs from naïve and 6-OHDA lesioned mice, the percent reduction in EPSC amplitude produced by Ro 25-6981 was roughly 25% (Fig. 2d, e, f); however, in iSPNs from dyskinetic mice, the contribution of GluN2B-containing NMDARs was about 40% (Fig. 2d, e). This change was similar in mice treated with levodopa for 3–5 or 12–14 days (Fig. 2f).

**Figure 2:**
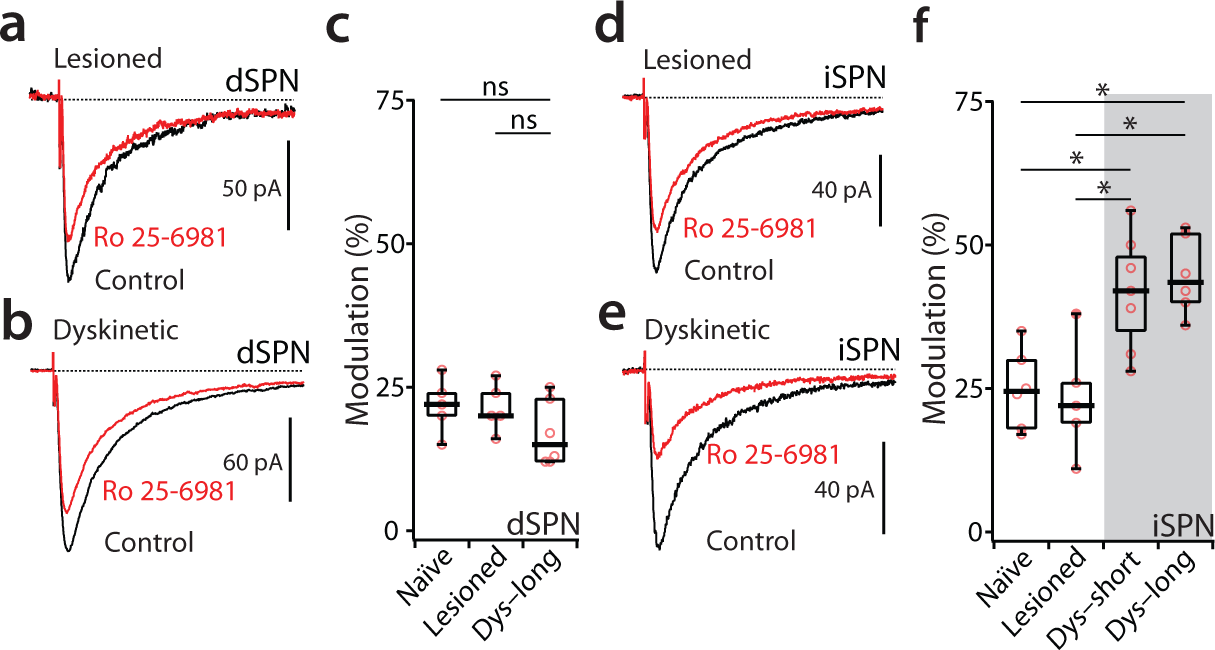
GluN2B-preferring inhibitor suppresses more NMDAR currents in levodopa-treated iSPNs than untreated iSPNs. (**a** and **b**) NMDAR currents were measured at the holding potential of –40 mV in the presence of 10 µM AMPAR antagonist NBQX and 10 µM GABA_A_ antagonist gabazine. In dSPNs, bath application of GluN2B selective inhibitor Ro 25-6891 (1 µM) reduced NMDAR currents by a similar size from naïve, lesioned and dyskinetic mice. (**c**) Box plot summary of the percent modulation of dSPN NMDAR currents from naïve (n=5, from 4 mice), lesioned (n = 5, from 3 mice) and dyskinetic (n = 6, from 4 mice). p > 0.05, Mann-Whitney test. ns not significant. (**d** and **e**) In iSPNs, Ro 25-6891 had more percent reduction in NMDAR currents from dyskinetic mice than naïve and lesioned mice. (**f**) Box plot sum of the percent modulation of iSPN NMDAR currents from naïve (n = 6, from 4 mice), lesioned (n = 5, from 4 mice), dyskinetic short-duration (3-5 days) (n = 7, from 5 mice) and dyskinetic long-duration (12-14 days) (n = 5, from 4 mice). * p < 0.05, Wilcoxon test.

Taken together, these results show that with LID induction, there is an up-regulation in GluN2B-containing NMDARs at iSPN glutamatergic synapses, but not at dSPN glutamatergic synapses. Given that in naïve iSPNs synaptic NMDARs appear to be largely (if not entirely) GluN2B-containing (based upon the P_o_ estimates), the up-regulation in the contribution of GluN2B subunits in the dyskinetic state could reflect an increased abundance of di-heteromeric GluN1/GluN2B NMDARs at iSPN glutamatergic synapses.

### LID induction was correlated with an increased abundance of iSPN silent synapses

Following 6-OHDA lesioning, spine density in iSPNs falls by roughly a third, presumably reflecting a form of homeostatic plasticity that accompanies the loss of inhibitory D2R signaling ^13^. With repeated treatment at dyskinesiogenic doses of levodopa, iSPN spine density returns into a normal range when measured after levodopa has cleared (off-state) ^13, 15, 25^. Thus, it is possible that the up-regulation in the contribution of GluN2B-containing NMDARs to evoked currents following LID induction reflects the addition of new axospinous synapses, rather than a modification of pre-existing synapses. In the developing brain, many glutamatergic synapses are dominated by GluN2B di-heteromeric NMDARs and have relatively few AMPARs, leading to ‘silent’ synapses that don’t pass current at resting membrane potentials (∼−70 mV) where NMDARs are blocked by Mg^2+^ ^32, 45^.

To determine whether SPNs up-regulate AMPA-deficient, silent synapses in the dyskinetic state, two different assays were employed. The first approach used minimal local stimulation to activate a small, random population of glutamatergic synapses and whole-cell voltage clamp methods to monitor postsynaptic currents. Given the stochastic nature of synaptic transmission, each round of stimulation in this protocol will evoke glutamate release at some synapses, but not others. For the postsynaptic cell to detect release, glutamate receptors must be present to generate detectable currents. If some of the synapses lack AMPARs but do have NMDARs, then the frequency of failures (where there is no detectable evoked current) will be voltage-dependent: that is, at membrane potentials where NMDARs are blocked by Mg^2+^ (∼−70 mV) the failure rate will be higher than at membrane potentials where NMDARs can report glutamate release (∼+40 mV). If it is assumed that the presynaptic release sites are independent and the release probabilities are the same at all the synapses, then the percentage of silent synapses can be estimated using the equation: 1 – [ln(F_–70_) / ln(F_+40_)], where F_–70_ is the failure rate at −70 mV and F_+40_ is the failure rate at +40 mV ^46^. This assay suggests that the percentage of dSPN silent synapses was not discernibly different in 6-OHDA lesioned and LID mice (∼ 20%, Fig. 3a-c). However, in iSPNs from LID mice the percentage of silent synapses was significantly higher after a 3-5 days or 12-14 days treatment of levodopa (∼40-50%, Fig. 3d-f).

**Figure 3:**
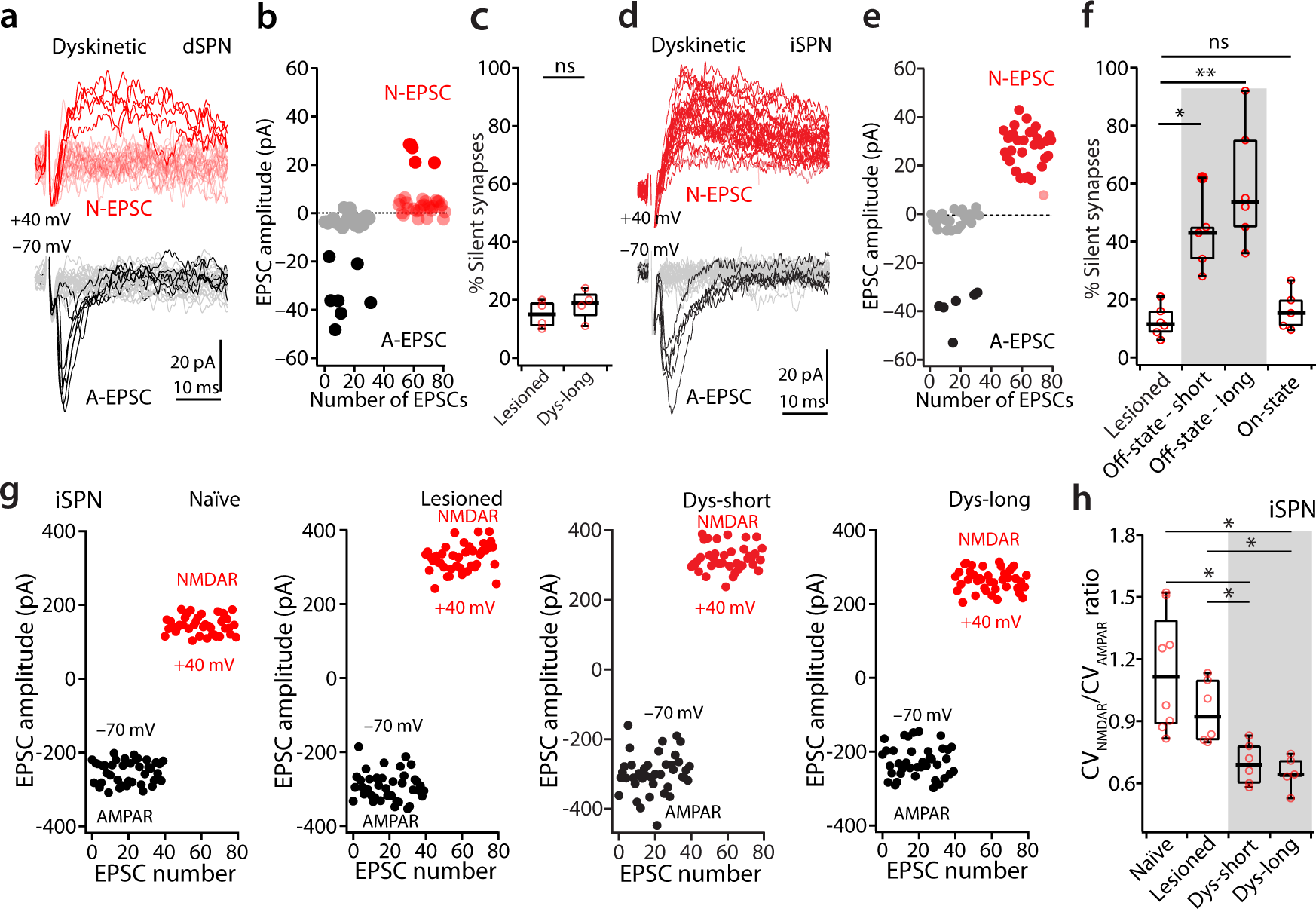
Enrichment of GluN2B-containing NMDARs creates AMPA-deficient silent synapses on iSPNs. Representative traces showing 40 consecutive EPSCs at +40 mV and – 70 mV from the levodopa-treated dSPN (**a**) and iSPN (**d**). The EPSCs were elicited by minimal electrical stimulation measured at –70 mV (AMPAR-EPSC) and +40 mV (NMDAR EPSC) in the presence of the AMPAR antagonist NBQX (10 µM). Plots of AMPAR and NMDAR EPSC amplitudes in the example dSPN (**b**) and iSPN (**e**). (**c**) Box plot sum of dSPN silent synapse percentage from lesioned (n = 4, from 3 mice) and dyskinetic states (n = 4, from 3 mice). The percent of silent synapses was estimated using the equation: 1 – [ln(F_–70_) / ln(F_+40_)], where F_–70_ is the failure rate at −70 mV and F_+40_ is the failure rate at +40 mV. ns not significant, Wilcoxon test. (**f**) Box plot summary of the percentage of iSPN silent synapses measured from lesioned (n = 6, from 5 mice), dyskinetic on-state (n = 5, from 5 mice), dyskinetic off-state short-duration treated (3-5 days, n = 5, from 4 mice) and dyskinetic off-state long-duration treated (12-14 days, n = 6, from 5 mice) animals. * p < 0.05, ** p < 0.01, Wilcoxon test. (**g**) Plots of iSPN AMPAR (at –70 mV) and NMDAR (at +40 mV) EPSCs from naïve, lesioned, dyskinetic (short duration) and dyskinetic (long-duration) conditions. (**h**) Box plot summary of the ratio of CV_NMDAR_ to CV_AMPAR_ from naïve (n = 7, from 5 mice), lesioned (n = 6, from 5 mice), dyskinetic short (n = 6, from 5 mice) and dyskinetic long (n = 5, from 5 mice) states. * p < 0.05, Wilcoxon test.

These measurements were all taken in mice well after the last dose of levodopa, mimicking the off-state. To assess whether this change was state-dependent, the minimal local stimulation experiments were repeated in *ex vivo* brain slices taken from dyskinetic mice an hour after levodopa treatment. Surprisingly, in on-state iSPNs, the relative abundance of silent synapses was no different than that in levodopa-naïve, 6-OHDA lesioned mice and significantly less than that in the off-state (Fig. 3f). Hence, the change in iSPN synaptic function was state-dependent.

To complement the failure rate assay, the relative variability of EPSCs was examined. In general, the coefficient of variation (CV) of EPSCs is inversely related to the number of contributing synapses. If there were more silent synapses, then the CV of NMDAR EPSCs (CV_NMDAR_) should be lower than that of AMPAR EPSCs (CV_AMPAR_). To test this hypothesis, the CV of NMDAR EPSCs and AMPAR EPSCs was measured. As expected, after either a short- or long-term treatment of levodopa, the CV_NMDAR_ / CV_AMPAR_ ratio fell in iSPNs (Fig. 3g, h).

### Up-regulation in GluN2B NMDARs enhanced LTP in iSPNs

In iSPNs, the induction of spike-timing dependent LTP relies upon the coordinated activation of NMDARs and A2a-Rs ^11, 47, 48^. Given their well-established linkage to LTP induction ^49, 50^, the up-regulation in GluN2B-containing NMDARs in iSPNs from dyskinetic mice should promote LTP induction, particularly in the LID off-state when D2R signaling is low and A2aR signaling is nominally high ^51–54^. To test this hypothesis, iSPNs from off-state LID mice were subjected to perforated patch, current-clamp recording in *ex vivo* brain slices and a pre-post, spike timing dependent plasticity (STDP) induction protocol was applied (Fig. 4a, b). In this protocol, afferent fibers were stimulated in short bursts (3 stimuli) with a theta electrode placed within approximately 100 microns from the cell body and trailing back-propagating action potentials (bAPs) (delay 5 ms) generated by short (1000 msec), intrasomatic current pulses. The pairing protocol was repeated every 200 msec for 2 seconds. This pairing protocol resulted in a robust LTP of glutamatergic EPSPs in naïve iSPNs (Fig. 4c, d). In agreement with earlier work ^9^, in iSPNs from levodopa-naïve, 6-OHDA lesioned mice, this protocol failed to produce any significant change in EPSP amplitude (Fig. 4d). However, STDP LTP was induced in iSPNs from LID mice subjected to short-term (3-5 days) (Fig. 4d) or long-term (12-14 days) (Supplementary Figure 1) systemic administration of levodopa.

**Figure 4:**
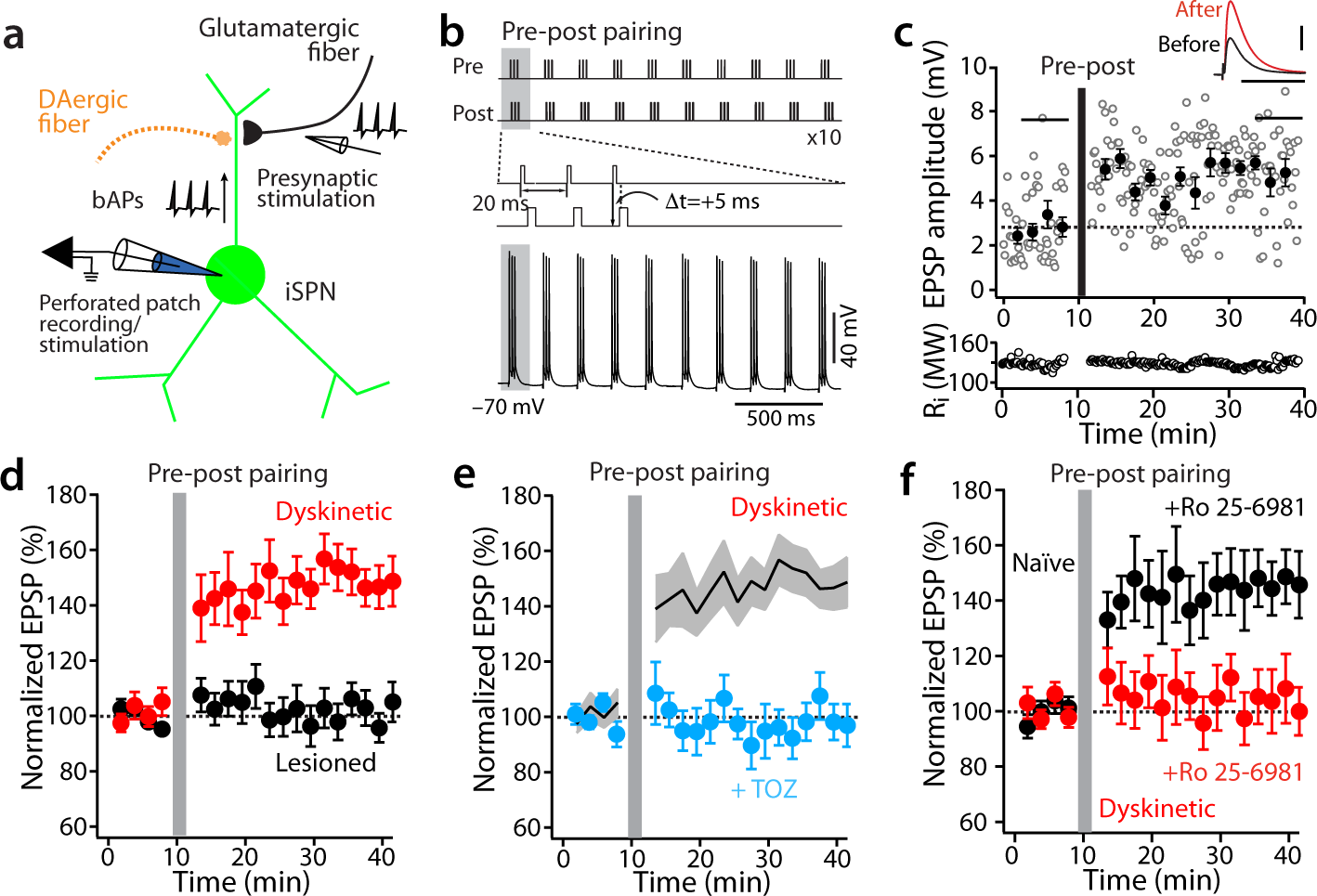
Up-regulation of iSPN GluN2B-containing NMDARs promotes LTP. (**a**) The experimental configuration. (**b**) Schematic illustrating the LTP induction protocol. (**c**) LTP was induced by a pre-post timing pairing in iSPNs. Plots show EPSP amplitude and input resistance (R_i_) as a function of time. Scale bars: 2 mV x 50 ms. (**d**) In the lesioned state, a pre-post timing pairing did not lead to LTP induction, whereas the same protocol revealed LTP in the dyskinetic state. Data are represented as mean ± SEM. Plot of the average EPSP amplitudes as a function of time. Lesioned n = 6, from 6 mice; dyskinetic n = 6, from 6 mice, p <0.05, Mann-Whitney rank sum test. (**e**) In dyskinetic state, pre-post pairing led to LTP; the LTP was disrupted by addition of the selective A2aR antagonist tozadenant (1 µM). Data are represented as mean ± SEM. Plot of the average EPSP amplitudes as a function of time. Dyskinetic n = 6, from 6 mice; tozadenant n = 5, from 5 mice, p <0.05, Mann-Whitney test. (**f**) In naïve state, selective GluN2B antagonist Ro 25-6891 (1 µM) did not disrupt LTP induction. However, the induction was blunted by the same inhibitor in the dyskinetic state. Data are represented as mean ± SEM. Plot of the average EPSP amplitudes as a function of time. Naïve n = 5, from 5 mice; dyskinetic n = 6, from 6 mice, p <0.05, Mann-Whitney test.

Pharmacological tools were used to assess the role A2aRs and GluN2B-containing NMDARs in the induction of STDP LTP. As expected, bath application of the A2aR antagonist tozadenant (1 µM) prevented LTP induction (Fig. 4e). Similarly, bath application of GluN2B antagonist Ro 25-6981 blunted STDP LTP induction in iSPNs from LID mice in the off-state (Fig. 4f). Interestingly, in brain slices from naïve mice, inhibiting GluN2B-containing NMDARs had no effect on pre-post STDP LTP induction (Fig. 4f), consistent with its modest effect on NMDAR currents in these cells (Fig. 1g).

### Off-state A2aR antagonism attenuated LID induction

In the healthy brain, the strength of iSPN glutamatergic synapses is widely thought to be shaped by the outcome of actions and linked activity in nigrostriatal dopaminergic neurons ^17, 55^ The strength of those connections governs the activity of iSPN ensembles and purposeful movement ^56^. In the brains of LID mice, where dopaminergic signaling is driven up and then down for hours by non-contingent, pulsatile levodopa administration, this relationship is lost. Our results show that this aberrant dopaminergic signaling up-regulates surface GluN2B-NMDARs in iSPNs and enables the induction of plasticity at glutamatergic synapses. This non-contingent plasticity could be a contributing factor in the emergence of the striatal pathophysiology underlying LID.

As a first step toward testing this hypothesis, an A2aR antagonist (tozadenant) with good brain bioavailability was given to 6-OHDA lesioned mice during levodopa treatment. If LTP induction in iSPNs was contributing to LID, then antagonizing A2aRs (which prevents LTP induction) should blunt LID induction. An important consideration in these experiments is the pharmacokinetic profile of tozadenant. Previous work has shown that co-administration of levodopa and the A2aR antagonist istradefylline (which has a similar pharmacokinetic profile as tozadenant) was ineffective in ameliorating LID in humans ^57^ as well as rodent models ^58^. However, our studies suggest that the critical time of A2aR signaling in the induction of iSPN synaptic plasticity is in the off-state, after levodopa has cleared, not in the on-state. To test this idea, mice were unilaterally lesioned with 6-OHDA and then randomly assigned to receive treatment with either levodopa (3 mg kg^−1^, i.p.) and vehicle or levodopa and tozadenant (30 mg kg^−1^, i.p.). Tozadenant was administered 5–6 hours after levodopa treatment, near the beginning of the off-state (Fig. 5a). AIM scores were recorded following the next dose of levodopa. As predicted, tozadenant significantly attenuated the development of AIMs over the treatment period (Fig. 5b, c). The reduction in dyskinesia scores did not occur at the expense of the anti-akinetic effect of levodopa, as forelimb use asymmetry was equally improved by levodopa when given alone or combined with tozadenant (Supplementary Fig. 2).

**Figure 5:**
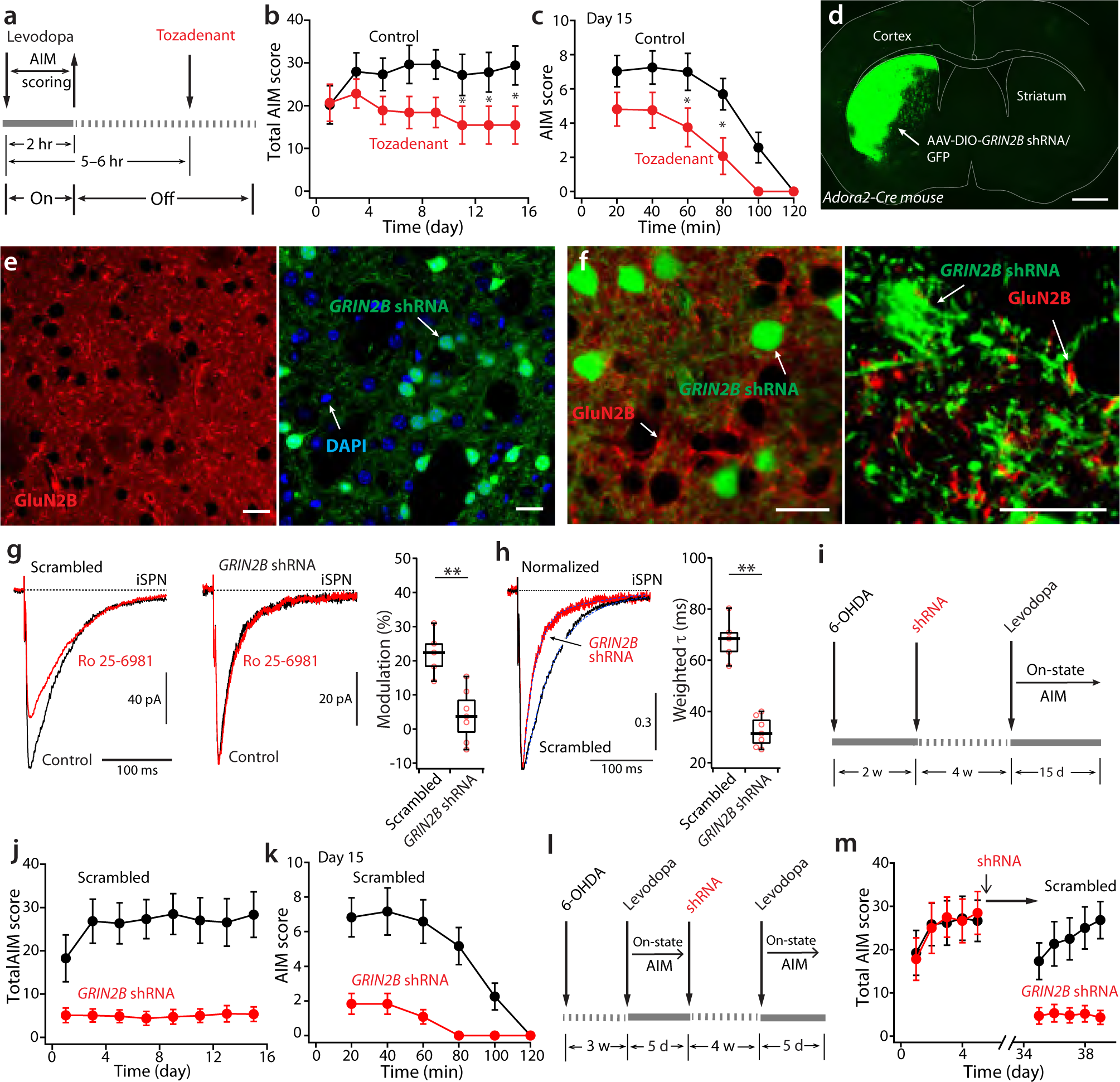
Antagonism of A2aR and GluN2B NMDARs attenuates the induction of dyskinetic behaviors. (**a**) Experimental timeline. Note that the treatment of tozadenant started in the off-state. (**b**) Plot of sum of axial, limb and orolingual AIM scores (levodopa + vehicle n = 8; levodopa + tozadenant n = 8, mean ± SEM) as a function of time. Systemic treatment with tozadenant (30 mg kg^−1^, i.p.) produced a significant overall reduction in AIM scores (time p = 0.0158, F(7, 98) = 2.62; group p = 0.114, F(1, 14) = 2.83; interaction p < 0.0001, F(7, 98) = 5.5; Fisher p < 0.05 on Day 11, 14 and 15. Repeated measure two-way ANOVA and post hoc Fisher). (**c**) Plot of sum of mouse AIM scores (n = 8, mean ± SEM) as a function of time on Day 15 (time p < 0.0001, F(5, 70) = 30.8; group p = 0.044, F(1, 14) = 4.88; interaction p = 0.0109, F(5, 70) = 1.8; Fisher p < 0.05 at 60 and 80 min. Repeated measure two-way ANOVA followed by post hoc Fisher). (**d**) Representative confocal image showing the expression of AAV-DIO-*GRIN2B* shRNA/GFP in the DLS of an *Adora2*-Cre mouse. Scale bar = 1mm. (**e, left**) Confocal image showing GluN2B immunoreactivity (red) in the contralateral striatum that did not have *GRIN2B shRNA*/GFP expression. Scale bar = 20 μm. (**e, right**) Confocal image showing *GRIN2B shRNA*/GFP (green) was expressed in ∼40% of total cells in the injected striatal region, consistent with Cre-dependent cell type specific expression in Adora2-Cre mice. Scale bar = 20 μm. (**f**) High-magnification merged image of GluN2B immunoreactivity (red) and *GRIN2B shRNA*/GFP (green). Note that the GluN2B immunoreactivity, which was concentrated in the dendrites, did not overlap at all with *GRIN2B shRNA*/GFP-positive dendrites. Scale bars = 20 μm. Knocking down GluN2B in infected iSPNs occluded selective antagonist blockade (**g**) and accelerated NMDAR current decaying (**h**). NMDAR currents were measured at the holding potential of –40 mV in the presence of 10 µM AMPAR antagonist NBQX and 10 µM GABA_A_ antagonist gabazine. (**g**) Box plot summary. Scrambled control n = 5, from 3 mice; *GRIN2B* shRNA n = 7, from 5 mice; p < 0.01, Mann-Whitney test. (**h**) Box plot summary. Scrambled control n = 5, from 3 mice; *GRIN2B* shRNA n = 7, from 5 mice; p < 0.01, Mann-Whitney test. (**i**) Experimental timeline. (**j** and **k**) Plots of sum of AIM scores (levodopa + scrambled shRNA (control) n = 6; levodopa + *GRIN2B* shRNA n = 6; mean ± SEM) as a function of days (**j**) or hours (**k**). (**j**) Time p <0.0001, F(7, 70) = 6.33; group p = 0.0021, F(1, 10) = 16.90; interaction p < 0.0001, F(7, 70) = 7.60; repeated measure two-way ANOVA and post hoc Bonferroni’s test. (**k**) Time p <0.0001, F(5, 50) = 22.47; group p = 0.001, F(1, 10) = 20.82; interaction p < 0.0001, F(5, 50) = 8.39; repeated measure two-way ANOVA followed by Bonferroni’s post test. (**l**) Experimental timeline. (**m**) Plot of sum of AIM scores (levodopa + scrambled shRNA (control) n = 6; levodopa + *GRIN2B* shRNA n = 6; mean ± SEM) as a function of time. (time p = 0.0009, F(4, 40) = 5.82; group p = 0.0045, F(1, 10) = 13.31; interaction p = 0.0005, F(4, 40) = 6.31; repeated measure two-way ANOVA followed by Bonferroni’s post test.).

### Knocking down the GluN2B subunit in iSPNs blunted both the induction and expression of LID

Although these experiments were consistent with our hypothesis, there was the possibility that the impact of systemically administered tozadenant on LID induction involved other cell types, like A2aR-expressing striatal cholinergic interneurons ^59^. To test the role of GluN2B up-regulation in iSPNs more directly, a genetic approach was used. A previously validated short hairpin ribonucleic acid (shRNA) sequence targeting *GRIN2B* mRNA ^60^ and a scrambled shRNA control were placed in a double-floxed inverse orientation (DIO) element and packaged in an adeno-associated virus (AAV) vector. In unilaterally 6-OHDA lesioned *Adora2*-Cre mice, the GluN2B-targeted AAV and its control were stereotaxically injected into the DLS (Fig. 5d). About 40% of the neurons in the striatal region injected expressed the FusionRed reporter, consistent with selective expression in iSPNs (Fig. 5e, f). To test for the efficacy of the knockdown, an antibody to GluN2B was used to examine the extent to which expression of the shRNA reduced expression of the protein. As expected, SPNs expressing the FusionRed reporter had little or no detectable GluN2B immunoreactivity (Fig. 5e, f). In addition, electrophysiological recordings found that *GRIN2B* knockdown in infected iSPNs occluded Ro 25-6981 sensitive current blockade and speeded up their de-activation (Fig. 5g, h). The intrinsic excitability of iSPNs (measured using intrasomatic current injection) was not altered by GluN2B knockdown in the LID off-state (Supplementary Fig. 3). Lastly, knocking down *GRIN2B* had no effect on the forelimb usage asymmetry produced by 6-OHDA lesioning (Supplementary Fig. 4)

Next, the impact of *GRIN2B* knockdown on dyskinetic behavior was examined. Four weeks after DLS injection of the AAV *GRIN2B* shRNA and its control, mice were given levodopa (3 mg kg^−1^, i.p.) daily for 15 days and LID AIM scores were rated every other day. Knocking down *GRIN2B* mRNA selectively in iSPNs significantly reduced peak AIM scores induced by levodopa and shortened the duration of dyskinetic behaviors (Fig. 5i-k). The reduction in dyskinesia scores did not occur at the expense of the anti-parkinsonian effects of levodopa, as forelimb use asymmetry was improved by levodopa in mice injected with either *GRIN2B* shRNA or its scrambled control (lesioned n = 6 mice; levodopa + *GRIN2B* shRNA n = 6 mice, p < 0.01, Wilcoxon test; lesioned n = 6 mice; levodopa + *GRIN2B* shRNA scrambled n = 6 mice; p < 0.01, Wilcoxon test) (Supplementary Fig. 4).

The ability of iSPN-specific targeting of GluN2B to alleviate the induction of LID was encouraging, but the clinically more relevant question is whether the approach could reduce the expression of established dyskinetic behavior. To answer this question, a cohort of well-lesioned *Adora2*-Cre mice were given dyskinesiogenic doses of levodopa (3 mg kg^−1^, i.p.) for 5 days to establish a stable LID. Then, the *GRIN2B* shRNA and scrambled shRNA vectors were injected into the DLS. Four weeks later, mice were challenged with levodopa. In agreement with previous work ^61^, mice injected with the control shRNA AAV exhibited a robust dyskinesia in response to levodopa after the ‘drug holiday’. However, mice in which *GRIN2B* had been knocked down exhibited little or no dyskinesia in response to the levodopa challenge (Fig. 5l, m). The reduction in dyskinesia did not compromise the pro-movement effect of levodopa (lesioned n = 6 mice; levodopa + *GRIN2B* shRNA n = 6 mice, p < 0.01, Wilcoxon test; lesioned n = 6 mice; levodopa + *GRIN2B* shRNA scrambled n = 6 mice; p < 0.01, Wilcoxon test) (Supplementary Fig. 5). Thus, blunting the contribution of GluN2B subunits to iSPN NMDARs effectively reduced both the induction and expression of LID, without diminishing the benefits of levodopa.

## Discussion

Three conclusions can be drawn from the experiments presented. First, LID induction is accompanied by an up-regulation in synaptic GluN2-containing NMDARs (GluN2B-NMDARs) in off-state iSPNs, but not dSPNs. Second, this up-regulation is associated with increased ‘silent’ glutamatergic synapses and restoration of STDP LTP in iSPNs, a form of plasticity lost in DA-depleted, levodopa-naïve mice. Unlike normal STDP LTP in iSPNs, this LID-associated plasticity was blocked by inhibiting GluN2B-NMDARs. Third, knocking down *GRIN2B* expression and the GluN2B subunit specifically in iSPNs attenuated the induction and expression of LID without compromising the symptomatic benefit of levodopa.

### LID was correlated with the up-regulation in GluN2B-containing NMDARs specifically in iSPNs

Over the past several decades, a handful of studies have drawn a connection between LID and the up-regulation of striatal NMDARs ^22, 62–65^. These studies have typically relied upon biochemical approaches that do not take into account the profound heterogeneity of striatal SPNs, which respond in diametrically opposing ways to the elevation in DA driven by levodopa treatment. In part, this may account for a lack of concordance about LID-induced alterations in the synaptic localization of GluN2A- and GluN2B-containing NMDARs ^62, 66–70^. Another issue that has largely been ignored is the potential importance of whether tissue was taken from animals in the off-state or the on-state. Striatal NMDARs can rapidly traffic from intracellular compartments to the plasma membrane ^71, 72^, raising the possibility that levodopa-induced alterations in surface expression are state-dependent.

Our studies show that the induction of LID is accompanied by an up-regulation in synaptic GluN2B-NMDARs that was specific to iSPNs in the off-state. This up-regulation manifested itself in several ways. First, NMDAR-mediated currents in iSPNs became more sensitive to the GluN2B-NMDAR antagonist Ro-25-6981, and their deactivation slowed. In addition, the NMDAR/AMPAR ratio rose in iSPNs from LID mice. The change in the NMDAR/AMPAR ratio in iSPNs from LID mice appears to be at odds with previous work ^13^. However, this discrepancy is attributable to how synapses were activated. Here, axons were electrically stimulated to evoke a spike that propagated to the terminal to induce Ca^2+^ entry and glutamate release, whereas in the previous work, either terminals were optogenetically stimulated or glutamate was uncaged on spine heads. These latter approaches will likely increase postsynaptic glutamate concentration above that produced by a spike invading the abutting presynaptic terminal ^73^. This is important because AMPARs have a much lower affinity for glutamate, than do NMDARs ^31^. Thus, although both approaches provide meaningful information, electrical stimulation is more likely to provide a more biologically meaningful estimate of synaptic function. That said, electrical stimulation has drawbacks in this context, the most relevant of which is the inability to interrogate a specific type of synapse. DLS SPNs are innervated by a variety of cortical and thalamic glutamatergic neurons ^74^. Going forward, optogenetic strategies could be used to evoke spikes in axons some distance from terminals, allowing an assessment of whether the up-regulation in GluN2B-NMDARs accompanying LID induction is input-specific.

Another apparent manifestation of GluN2B-NMDAR up-regulation in iSPNs was the growth in the number of ‘silent’ synapses – that is, synapses with NMDARs, but few or no AMPARs ^32, 45^. Again, this shift was specific to iSPNs. It also was specific to the off-state, as the number of silent synapses returned to control values in the on-state. This cellular and state specificity could have its basis in the changing modulator environment associated with on- and off-states. NMDAR trafficking to the surface is known to be enhanced by phosphorylation of GluN2B at Tyr 1472 by Fyn kinase, a member of the src kinase family ^75,76^. Fyn kinase is anchored at glutamatergic synapses by PSD95 and is positively modulated by G-protein coupled receptors (GPCRs) that stimulate adenylyl cyclase (AC) ^77^. Previous work has shown that striatal D1Rs increase Fyn kinase phosphorylation of Tyr 1472 and promote synaptic localization of GluN2B-NMDARs ^78^. In contrast, D2Rs, which are expressed by iSPNs, inhibit AC and reduce Fyn kinase activity ^79^. In iSPNs, D2Rs are key negative regulators of A2aR-mediated stimulation of AC ^80^. Thus, in the off-state, when D2R signaling is low and constitutive A2aR signaling is dis-inhibited, Fyn kinase activity should rise, leading to phosphorylation of GluN2B-NMDARs and increased surface expression. Consistent with this hypothesis, phosphorylated Fyn kinase is elevated in the striatum of dyskinetic rodents sacrificed in the off-state ^81^. In the on-state, the sustained activation of iSPN D2Rs should suppress Fyn kinase activity and the trafficking of GluN2B-NMDARs to the surface – accounting for the normalization of silent synapse numbers in the on-state.

### In the off-state, LTP in iSPNs was enabled by GluN2B-NMDARs

Following lesioning of the nigrostriatal dopaminergic system in rodents, corticostriatal synaptic plasticity in SPNs is lost in the ensuing weeks ^7–9^. Subsequent levodopa treatment restores STDP LTP in D1R-expressing dSPNs ^9^. However, as D1R signaling with levodopa treatment is sustained, the resulting plasticity is disconnected from normal movement control and contributes to the induction of dyskinesia ^4^. Indeed, counter-acting D1R coupling to AC during the on-state by enhancing M4 muscarinic receptor (M4R) activity diminishes LID induction ^9^.

Our studies have identified a complementary plasticity in iSPNs occurring in the off-state. In *ex vivo* brain slices from dyskinetic mice in the off-state, a conventional pre-post pairing protocol was able to induce a robust STDP LTP in iSPNs recorded in perforated patch mode. The same protocol was not able to induce LTP in iSPNs from 6-OHDA lesioned mice that had not been treated with levodopa. As in iSPNs from normal healthy mice, the STDP LTP in iSPNs was dependent upon activation of A2aRs and NMDARs. However, unlike the situation in naïve iSPNs, the STDP in iSPNs taken from dyskinetic mice was disrupted by antagonizing GluN2B-NMDARs. Based upon work elsewhere ^82, 83^, it is tempting to speculate that the off-state STDP LTP in iSPNs largely reflects the trafficking of AMPARs into silent GluN2B-enriched glutamatergic synapses. Regardless, the persistent, non-contingent elevation in the ability to induce LTP in the off-state is likely to result in the distortion of synaptic weight sculpted by experience, contributing to the aberrant movement control characteristic of LID.

### Disrupting GluN2B-NMDAR function in iSPNs blunted both the induction and expression of LID

Although the up-regulation in GluN2B-NMDAR expression and restoration of LTP in off-state iSPNs were correlated with LID, it is possible they were consequences, not causes ^5^. To distinguish between these possibilities, the impact of two interventions on LID behaviors were examined. First, the A2aR antagonist tozadenant was given to 6-OHDA lesioned mice after the pro-motoric effects of levodopa had waned, at the beginning of the off-state. Interestingly, this treatment blunted the induction of LID. This observation is not at odds with previous reports that a similar A2aR antagonist given concurrently with levodopa had no effect because of the different temporal profile of target engagement ^58, 84, 85^. The most parsimonious explanation for the anti-dyskinetic effects of off-state A2aR antagonism is that it effectively impeded GluN2B-NMDAR trafficking to the surface and the induction of aberrant LTP. That said, there are other potential explanations, like disruption of A2aR signaling in striatal cholinergic interneurons ^59^.

The more definitive experiment was the genetic one: knocking down *GRIN2B* expression specifically in DLS iSPNs. These experiments took advantage of *Adora2*-Cre mice and AAV delivery of a Cre-dependent *GRIN2B* shRNA construct. The efficacy of this knockdown was confirmed by both immunocytochemical and electrophysiological assays. *GRIN2B* knockdown in iSPNs dramatically reduced the induction of LID. But it also did something that was more unusual. When the knockdown was performed after establishing LID, it prevented the expression of dyskinesia when levodopa treatment was re-instated without compromising its motor benefits.

Largely in agreement with previous studies ^4, 16^, our work suggest that the striatal pathophysiology underlying LID involves both cell-type and state-specific adaptations triggered by the oscillation in brain DA concentration following levodopa administration. There is no doubt that during the on-state, the sustained elevation in striatal DA levels drives dSPN hyper-activity, which is critical to dyskinetic movement ^4, 5, 9, 14^. However, striatal movement control is mediated by an interaction between the circuitry anchored by both dSPNs and iSPNs. Inappropriately timed or scaled activity in either circuit can disrupt normal movement. Our results show that rather than being a passive period following episodes of dyskinesia, the off-state is a period of active remodeling in iSPNs that is triggered by the profound imbalance in neuromodulatory signaling. This GluN2B-dependent remodeling appears to be essential for LID expression hours later when levodopa is re-administered. Precisely why this is the case remains to be elucidated. Recent work has highlighted the importance of reduced iSPN excitability during the on-state in LID ^16, 86^. The engagement of GluN2B-containing NMDARs in iSPNs during the off-state could promote subsequent on-state hypoexcitability by triggering homeostatic mechanisms that promote long-term depression of excitatory synaptic inputs and depress intrinsic excitability ^87^.

Indeed, both iSPN intrinsic excitability and corticostriatal synaptic strength fall during the on-state [unpublished observations]. Whether these on-state adaptations are blunted by knocking down *GRIN2B* expression remains to be determined. Alternatively, GluN2B deletion might prevent the off-state alterations in iSPN glutamatergic synapses that drive inappropriately timed or scaled activity and unwanted movement in the on-state. Aberrant activity in iSPN ensembles also could disrupt dSPN ensembles regulated by both short and long iSPN feedback loops ^88, 89^.

In addition to revealing a new facet of abnormal striatal plasticity in LID, these studies point to the potential of iSPN-targeted GluN2B-reducing strategies for alleviating established LID. The lack of clinical success in previous attempts to globally reduce GluN2B-NMDAR function with systemically administered drugs is likely to stem from limitations posed by off-target effects. The rapid advancement of therapies that yield cell-type specific gene expression with systemically administered vectors ^90, 91^ raises the hope that a novel treatment for LID is on the horizon.

## Acknowledgments

This work was supported by NS 34696 and the JPB Foundation.

## Methods

### Animals

Male C57Bl/6 mice expressing tdTomato or eGFP under control of either the *Drd1a* or *Drd2* receptor regulatory elements (NINDS GENSAT BAC Transgenics Project, Rockefeller University, New York, NY) were used. All the mice used for the study were hemizygous for these transgenes. BAC transgenic mice expressing Cre recombinase under control of the A2aR regulatory elements were used for *GRIN2B* shRNA experiments. All mice were 8–10 weeks of age before stereotaxic surgery. Animal use procedures were reviewed and approved by the Northwestern Institutional Animal Care and Use Committee.

### Slice preparation and electrophysiology

Mice were deeply anesthetized intraperitoneally with a mixture of ketamine (50 mg kg^−1^) and xylazine (4 mg kg^−1^) and perfused transcardially with 5–10 ml of ice-cold artificial CSF (aCSF) comprising (in mM): 125 NaCl, 2.5 KCl, 1.25 NaH_2_PO_4_, 2.0 CaCl_2_, 1.0 MgCl_2_, 26 NaHCO_3_ and 10 glucose (305 mOsm l^−1^). Parasagittal slices were cut in ice-cold external solution containing (mM) 110 choline chloride, 26 NaHCO_3_, 1.25 NaH_2_PO_4_, 2.5 KCl, 0.5 CaCl_2_, 7 MgCl_2_, 11.6 sodium ascorbate, 3.1 sodium pyruvate and 5 glucose (305 mOsm l^−1^).

Experiments were performed in the dorsolateral striatum at elevated temperature (30–31°C). Patch pipettes were loaded with internal solution containing (mM): 120 CsMeSO_3_, 15 CsCl, 8 NaCl, 10 HEPES, 0.2 EGTA, 10 TEA Chloride, 5 QX-314, 2 Mg-ATP, 0.3 Na-GTP (305 mOsm l^−1^) for whole-cell voltage-clamp recordings; or 115 K-gluconate, 20 KCl, 1.5 MgCl_2_, 5 HEPES, 0.2 EGTA, 2 Mg-ATP, 0.5 Na-GTP, 10 Na-phosphocreatine (280 mOsm l^−1^) for whole-cell current-clamp recordings. All the recordings were made using a MultiClamp 700B amplifier, and signals were filtered at 2 kHz and digitized at 10 kHz. Data were discarded when the series resistance (voltage-clamp) or input resistance (current-clamp) changed >20% over the time course of the experiment. For perforated patch recordings, the internal recording solution comprised (in mM): 125 KMeSO_4_, 14 KCl, 2 MgCl_2_, 0.2 EGTA, 10 HEPES. Amphotericin B was used to achieve electrical access through the perforated-patch method ^9, 27^.

Long-lasting synaptic plasticity was induced using protocols consisting of subthreshold synaptic stimulation paired with somatically induced action potentials (APs) at theta frequency (5 Hz). These protocols consisted of 10 trains of ten bursts repeated at 0.1 Hz, with each burst composed of three APs preceded with three EPSPs at 50 Hz (pre-post timing pairing, +5ms). To ensure induction of consistent synaptic plasticity, postsynaptic neurons were depolarized to –70 mV from their typical resting membrane potentials (–85 mV) during their induction. GABA_A_ receptor was blocked by the bath application of gabazine (10 µM).

### Mouse unilateral 6-OHDA model and LID

Mice were anaesthetized with an isoflurane precision vaporizer, placed in a stereotaxic frame, and a hole was drilled over the MFB. After exposing the skull, 3.5 mg ml^−1^ free base 6-OHDA hydrochloride with 0.02% ascorbic acid was injected using a calibrated glass micropipette at the following coordinates: AP: –0.7; ML: –1.2; DV: –4.75. Two-three weeks after surgery, the degree of damage to nigrostriatal DA neurons was assessed with a forelimb-use asymmetry test ^9, 26, 27^. Well-lesioned animals were then assigned to receive either *GRIN2B* shRNA or its scrambled control in the DLS in a total of 6 sites at the following coordinates: AP: 0.9; ML: –2.3; DV: –3.4; −2.8; AP: 0.6; ML: –1.5; DV: –3.4; −2.8; AP: 0.24; ML: –1.9; DV: –3.3; −2.7.

One day after the forelimb-use asymmetry test, mice underwent behavioral testing for AIMs following levodopa treatment as previously described ^9, 27, 92^. For *ex vivo* brain slice recording, animals received daily levodopa for either short (3–5 days) or prolonged period (12–14 days). Benserazide was co-administered at 12 mg kg^−1^ to inhibit peripheral conversion of levodopa to DA. AIMs (axial, limb and orolingual movements) were rated as previously described ^93^. Each animal was observed individually for 1 min every 20 min for 2 hours. Physiological experiments were performed one day (off-state) or one hour (on-state) after the last levodopa administration.

### Immunostaining and confocal microscopy

After behavioral experiments, mice that had been injected with AAV-DIO-*GRIN2B* shRNA/GFP were anesthetized as described above and perfused transcardially with saline for ∼1 min and then with ice-cold 4% paraformaldehyde (wt/vol) in 1x phosphate buffered saline (4% PFA-PBS). The mouse brains were dissected out, post-fixed in 4% PFA-PBS overnight at 4°C, and sectioned into coronal slices (100 μm-thick) using a Leica vibratome (VT1200, Leica). Immunostaining was performed using methods described previously ^94^. Briefly, the fixed brain slices were permeabilized and blocked in PBS containing 5% normal goat serum and 0.2% Triton-X100 (NGS-PBST) for 1 hr at 4°C, and then incubated with mouse monoclonal anti-NMDAR2B antibody (1:100 dilution in NGS-PBST, ab93610, Abcam) overnight at 4°C. After four washes in NGS-PBST, the slices were incubated with goat anti-mouse Alexa Fluor 555 secondary antibody (1:1000 dilution in NGS-PBST, A-21422, Invitrogen) for 2 hours at room temperature. Slices were washed with NGS-PBST four times, counter-stained with DAPI (1:1000 dilution in PBS, D1306, Invitrogen), and washed again with PBS once. Slices were then mounted with VECTASHIELD Mounting Medium (Vector Laboratories) and imaged under a laser scanning confocal microscope (FV10i-DUC; Olympus). Images were adjusted for brightness, contrast, and pseudo-coloring in ImageJ (US National Institutes of Health).

### Viral vectors

The shRNA sequence targeting *GRIN2B* mRNA ^60^ was inserted into the miR30 backbone of LENG plasmid, a gift from Johannes Zuber (Addgene plasmid # 111162) ^95^ and then into 3’-UTR of GFP in pAAV-Ef1a-DIO GFP plasmid, a gift from Karl Deisseroth (Addgene plasmid # 27056). YFP was substituted for GFP in this construct. The hairpin for the miR30-based expression of scrambled shRNA, a gift from William Wisden (Addgene plasmid #71383) ^96^ was inserted into 3’-UTR of GFP in pAAV-Ef1a-DIO GFP plasmid in similar way. Viruses were packaged by Virovek (https://www.virovek.com/).

### Data analysis and statistics methods

Data analysis was conducted with Igor Pro 8 and Clampfit 10. EPSP amplitude was calculated from 50 sweeps immediately before the start of induction and 20–30 min after the end of induction. Compiled data were expressed as mean ± SEM. Statistical tests were performed using Excel and SigmaStat. Nonparametric Mann-Whitney rank sum and Wilcoxon signed rank tests were used to assess the experiment results, using a probability threshold of 0.05.

Statistical analysis for behavioral data was carried out using Prism 10. Data were analyzed using parametric repeated measure two-way ANOVA followed by post hoc Fisher or Bonferroni’s test.

### Chemicals and reagents

Reagents were obtained as follows: amphotericin B, ZnCl_2_, tricine, 6-OHDA HCl, levodopa, benserazide HCl (Sigma); APV, NBQX, SR95531 (gabazine), MK-801, Ro 25-6891 (Tocris); tozadenant (APExBIO).

### Study approval

Present studies in animal use were reviewed and approved by the Northwestern Institutional Animal Care and Use Committee (Chicago, IL).

---

**Supplementary Figure 1:**
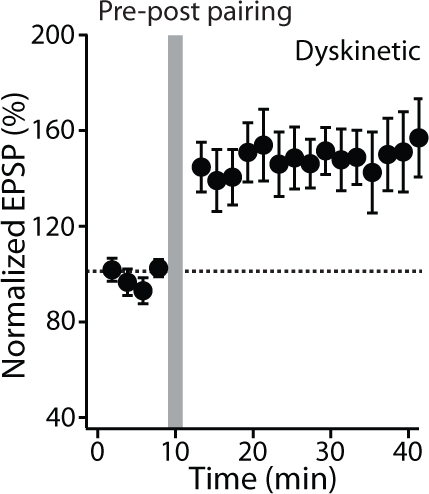
Prolonged treatment (12-14 days) of levodopa promotes induction of iSPN LTP in the LID off-state. LTP was induced with the pre-post pairing protocol in iSPNs from levodopa treated lesioned animals (n = 5; p < 0.05, Wilcoxon test). Plot of normalized EPSP amplitude as a function of time. Data are shown as mean ± SEM. The dashed line represents the average of EPSP amplitude before induction. The vertical bar indicates STDP induction.

**Supplementary Figure 2:**
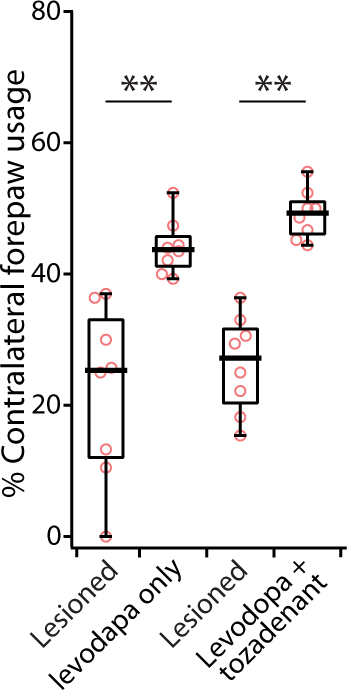
Co-treatment of the A2aR antagonist tozadenant does not jeopardize the anti-parkinsonian effect of levodopa. The reduction in AIM scores had no effect on the anti-parkinsonian benefit of levodopa, as forelimb use asymmetry was improved by levodopa only or when co-treated with tozadenant (levodopa only n = 8 mice, p < 0.01, Wilcoxon test; levodopa + tozadenant n = 8 mice, p < 0.01, Wilcoxon test). ** p < 0.01.

**Supplementary Figure 3:**
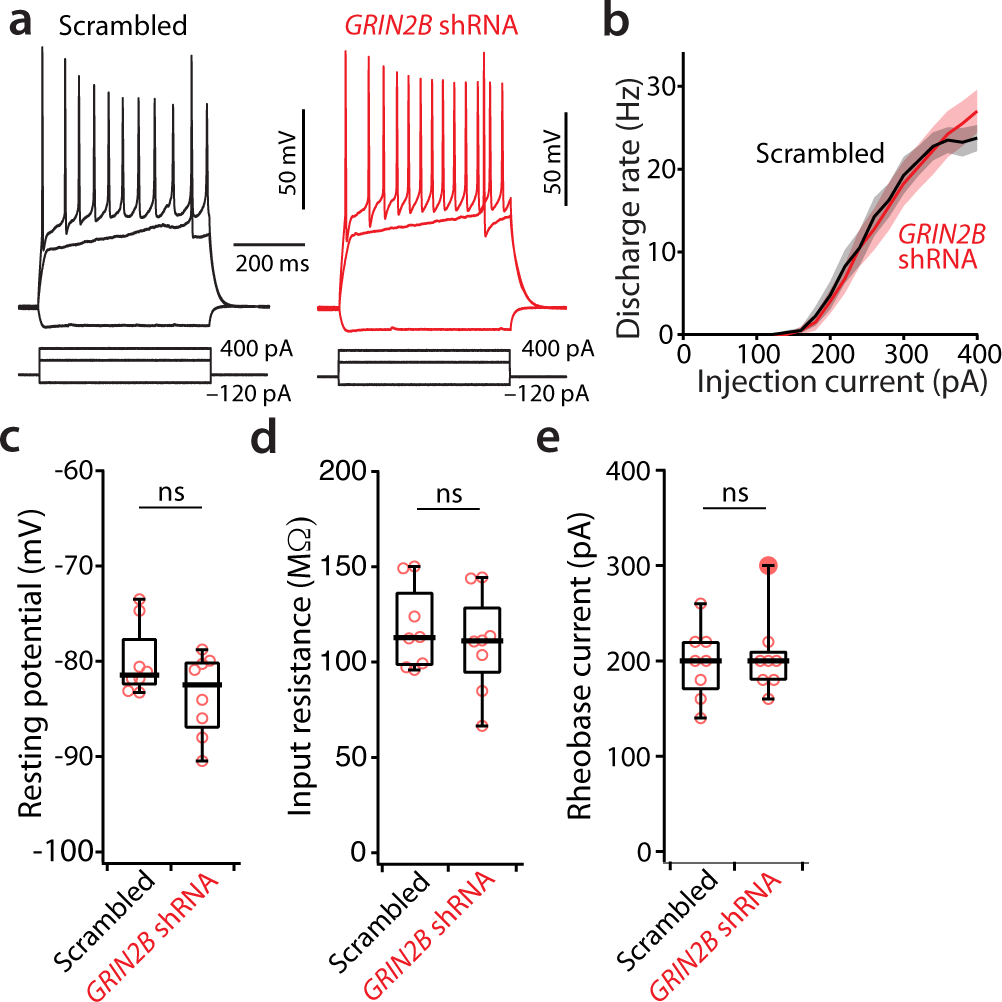
iSPN intrinsic properties were not altered with GluN2B knockdown in LID off-state. (**a**) representative current clamp recording traces showing the change of membrane potentials and evoked action potentials in response to –120 pA, 220 pA and 400 pA current injection steps. Left, scrambled control (black); right, *GRIN2B* shRNA (red). (**b**) The firing induced by a series of 0-400 pA current injection steps was not changed with *GRIN2B* shRNA (top middle; n = 8 cells/3 mice for each group). There is no difference in (**c**) resting membrane potential (scrambled control: median = –81.4 mV; *GRIN2B* shRNA: median = –82.5 mV; ns not significant), (**d**) input resistance measured with a –20 pA hyperpolarizing step (scrambled control: median = 112.8 MΩ; *GRIN2B* shRNA: median = 111.1 MΩ; ns not significant), and (**e**) rheobase current — current required to evoke first action potential (scrambled control: median = 200 pA; *GRIN2B* shRNA: median = 200 pA; ns not significant) between scrambled control and *GRIN2B* shRNA groups.

**Supplementary Figure 4:**
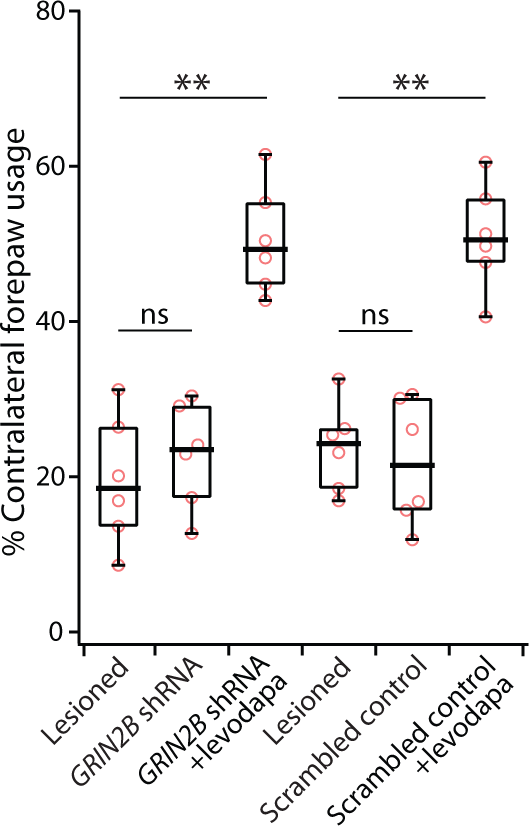
Knocking down *GRIN2B* has no effect on the forelimb use asymmetry elicited by 6-OHDA and does not imperil the symptomatic benefit of levodopa. Down-regulation of GluN2B containing NMDARs in iSPNs did not change the forelimb use asymmetry produced by 6-OHDA lesioning (*GRIN2B* shRNA n = 6 mice; p > 0.05, Wilcoxon test; scrambled control n = 6 mice; p > 0.05, Wilcoxon test). The reduction in AIM scores had no effect on the anti-parkinsonian benefit of levodopa, as the forelimb use asymmetry was improved by levodopa and *GRIN2B* shRNA or its scrambled control (levodopa + *GRIN2B* shRNA n = 6 mice, p < 0.01, Wilcoxon test; levodopa + scrambled control n = 6 mice, p < 0.01, Wilcoxon test). ** p < 0.01, ns not significant.

**Supplementary Figure 5:**
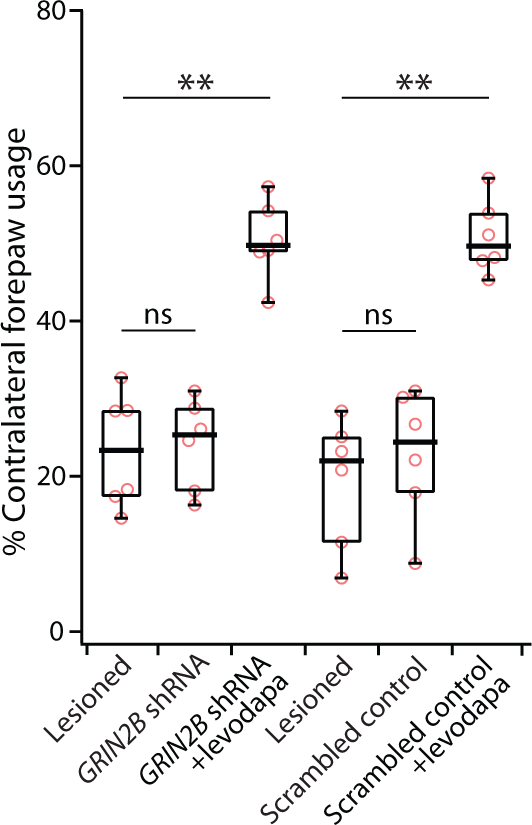
Down-regulation of *GRIN2B* has no effect on the forelimb use asymmetry elicited by 6-OHDA and does not compromise the anti-parkinsonian benefit of levodopa. Knock-down of iSPN *GRIN2B* did not change the forelimb use asymmetry produced by 6-OHDA lesioning (*GRIN2B* shRNA n = 6 mice; p > 0.05, Wilcoxon test; scrambled control n = 6 mice; p > 0.05, Wilcoxon test). The reduction in AIM scores had no effect on the anti-parkinsonian benefit of levodopa, as the forelimb use asymmetry was improved by levodopa and *GRIN2B* shRNA or its scrambled control (levodopa + *GRIN2B* shRNA n = 6 mice, p < 0.01, Wilcoxon test; levodopa + scrambled control n = 6 mice, p < 0.01, Wilcoxon test). ** p < 0.01, ns not significant.

